# Novel approach for carbon-wise utilization of lignin-related compounds by synergistically employing anaerobic and aerobic bacteria

**DOI:** 10.1101/2024.02.14.580265

**Authors:** Ella Meriläinen, Elena Efimova, Ville Santala, Suvi Santala

## Abstract

Lignin is a highly abundant but strongly underutilized natural resource that could serve as a sustainable feedstock to produce chemicals by microbial cell factories. However, the production from lignin-related aromatics is hindered by limited substrate range and inefficient catabolism of the production hosts. Particularly, the aerobic demethylation reactions are energy-limited and cause growth inhibition and loss of CO2. Here, we present a novel approach for carbon-wise utilization of lignin-related aromatics by the integration of anaerobic and aerobic metabolisms. In practice, we employed an acetogenic bacterium *Acetobacterium woodii* for anaerobic O-demethylation of aromatic compounds, which distinctively differs from the aerobic demethylation; in the process, the carbon from the methoxyl groups is fixated together with CO2 to form acetate while the aromatic ring remains unchanged. These accessible end-metabolites were then utilized by an aerobic bacterium *Acinetobacter baylyi* ADP1. Finally, we demonstrated the production of muconic acid from guaiacol, an abundant but inaccessible substrate to most microbes, with a nearly equimolar yield with only a minor genetic engineering and without the need for additional organic carbon source. This study highlights the power of synergistic integration of distinctive metabolic features of bacteria, thus unlocking new opportunities for harnessing microbial cocultures in upgrading challenging feedstocks.

## Introduction

Lignin is the second most abundant polymer in Nature, mainly consisting of three aromatic units comprising H- (*p-*coumaryl alcohol), G- (coniferyl alcohol), and S- (sinapyl alcohol) lignin types^1^. Large quantities of lignin, often considered as a waste material, are generated through pulping and biorefinery industries, agriculture, and wood processing^2^. Due to lignin’s complexity and heterogeneity, it is currently a strongly underutilized resource. However, it could serve as a renewable replacement for fossil-based resources in the production of chemicals and materials, for which biological lignin valorization is considered as the most promising approach. To access the monomers of the complex lignin polymer, the covalent C-C and C-O bonds connecting the units must be broken down via depolymerization or pyrolysis reactions^3^. The chemical structures of the obtained monomers are derivatives of the lignin units, varying with the number of their methoxyl and hydroxyl groups and the structure of the possible acrylate chain linked to the aromatic ring^4^.

Biological lignin valorization relies on the specific catabolic pathways of microbes, such as β- ketoadipate pathway^5^, to funnel heterogeneous mixtures of aromatic compounds into central carbon metabolites^6,7^. Previously several bacteria, including species of *Pseudomonas* and *Rhodococcus*, have been engineered to produce industrially relevant compounds, such as *cis,cis-*muconic acid (ccMA)^8–11^, β-ketoadipic acid and muconolactone^12^, as well as native products including polyhydroxyalkanoates (PHAs) and triacylglycerides (TAGs)^6^, from lignin-derived aromatic compounds. We have previously engineered *Acinetobacter baylyi* ADP1 (hereafter referred to as ADP1) to produce 1-alkenes or alkanes^13–15^ and wax esters^14^ from lignin-related substrates or technical lignin. In addition, ADP1 has also been utilized to produce mevalonate from lignin-related aromatics^16^.

However, the biological lignin valorization by microbes has its own hardships, including limited aromatic substrate range and the toxicity of the aromatic compounds causing growth inhibition at industrially relevant concentrations^6,17^. For example, the valorization of guaiacol (2-methoxyphenol), one of the main monomeric products obtained from alkaline-pretreated softwood lignin, has been set as an important goal^6^, but only few bacteria, like *Rhodococcus rhodochrous*^18^ and *Amycolatopsis sp.* ATCC 39116^19^ can naturally catabolize the compound. The ability to utilize guaiacol has previously been engineered in non-native hosts including *Pseudomonas putida* EM42^20^ and ADP1^21^. However, to achieve sufficient utilization of guaiacol, co-expression of redox partners^20^, or evolution-based approaches^21^ are often required.

In addition to the restricted substrate range, also the natural pathways exhibit inherent challenges: the G- and S-lignin derived compounds that contain one and two methoxyl groups, respectively, need to be demethylated prior to further catabolic steps. Aerobic demethylation is recognized as one of the major hurdles in the utilization of lignin-related aromatic compounds^22–24^. In aerobic conditions, O-demethylase reactions are catalyzed by enzymes belonging into either Rieske oxygenases (ROs), cytochrome P450 or tetrahydrofolate (THF)-dependent demethylases classes^25^. For example, VanAB enzyme in *P. putida* is a RO-type enzyme, which demethylates G-lignin-derivate vanillic acid^26^ and S- lignin-based syringic acid^27^ into protocatechuic acid (PCA) and gallic acid, respectively. Also ADP1 harbors similar VanAB system and it is known to convert G-lignin-derivates ferulic acid and vanillic acid into PCA^28^. However, the demethylation reactions catalyzed by ROs or P450s produce highly cytotoxic formaldehyde^29^ as their byproduct^22,26^. To eliminate formaldehyde, cells harbor detoxification routes^29^ which eventually degrade the compound into CO2 with a price of losing the carbon^23^. In addition, the aerobic demethylation reaction requires oxidation of NADH to NAD^+^, causing an intracellular cofactor imbalance and reduced primary metabolic activity^24^. Thus, to ensure efficient demethylation, a supporting carbon source maintaining the primary metabolism is often provided^24,30^. Because of these issues, novel approaches to circumvent the problems related to aerobic demethylation are required.

Acetogens are anaerobic bacteria that can grow autotrophically on H2-CO2 mixtures using Wood- Ljungdahl pathway for synthesizing acetyl-CoA from CO ^31,32^. Because of their ability to fix gaseous CO_2_, acetogens could play a vital role in biotechnology for the sustainable production of biochemicals. The usage of acetogens to provide organic carbon compounds from CO2 in two-stage processes was recently reviewed by Ricci *et al.*^33^: in this approach, the fermentation products of acetogens, such as acetate or ethanol, are subsequently utilized and upgraded by aerobic bacteria. Both acetate and ethanol enter the central metabolism as acetyl-CoA, making the number of possible end-products nearly limitless^33,34^. We have previously utilized two-stage processes where we first cultivated *Acetobacterium woodii* on CO2 to produce acetate, which was then used as a substrate by ADP1 for wax ester^35^ or alkane^36^ syntheses.

In addition to H2-CO2 mixtures, acetogens utilize a variety of organic C1-compounds^37^ as additional carbon sources and electron donors for the CO2 fixation. Interestingly, some acetogens can also utilize the methoxyl groups of aromatic compounds, including those derived from G-lignin, such as guaiacol, vanillic acid, and ferulic acid, by removing the methyl group by anaerobic O-demethylation.^38^ Demethylating reactions have been observed in several types of acetogens including *A. woodii*, *Moorella thermoacetica*, and *Sporumusa ovata*^39^. In *A. woodii,* the demethylation process involves the fixation of carbon from the methoxyl group and the simultaneous CO2 reduction into two-carbon molecule, acetate^40^. The biomass production and the molar yield of the produced acetate are positively correlated with the number of the methoxyl groups in the aromatic compound^41^. In addition to the methoxyl group demethylation, *A. woodii* is also able to reduce any double bonds located in the acrylate side chain of the aromatic compounds, and to use the bonds as electron acceptor instead of CO_2_ ^42^, which results in increased biomass but lower acetate production yields^41^. During these processes, the aromatic ring stays unchanged^41^.

In this study, we present a novel concept for upgrading lignin-related aromatic compounds into industrially interesting products while simultaneously overcoming the issues of aerobic demethylation and carbon loss. To achieve this, we integrate the best features of acetogens and aerobic bacteria. First, we investigated the suitability of the demethylation end products of *A. woodii* for *A. baylyi* ADP1 growth. Following this, we established a two-stage culture showing the compatibility of *A. woodii* and ADP1 for synergistic utilization of lignin-related aromatic compounds. Finally, as a proof-of-concept, we demonstrated an equimolar conversion of guaiacol, a model compound of G-lignin and an inaccessible substrate for ADP1, into MA by a one-pot coculture approach. In the process, *A. woodii* demethylates guaiacol into catechol, which is subsequently converted into MA by ADP1. Importantly, employing the anaerobic demethylation step ensured that the methoxyl group of guaiacol was not lost to CO2, but instead fixated into acetate, thus providing a carbon source for the growth of ADP1. Simultaneously, we were able to bypass the key bottleneck, aerobic demethylation, in the utilization of lignin-related aromatic compounds. This study highlights the potential of harnessing and integrating distinctive metabolic features of bacteria for improved carbon recovery and overcoming metabolic bottlenecks in the valorization of recalcitrant feedstock.

## Results

### Set-up for the utilization of aromatic compounds by integrating the metabolisms of anaerobic and aerobic bacteria

To investigate the utilization of lignin-related aromatic compounds by the collaboration of anaerobic and aerobic bacteria, we employed the acetogen *A. woodii* and the strictly aerobic ADP1 for the study. In the designed co-utilization scheme, *A. woodii* would utilize the methoxylated aromatic compounds via demethylation reaction, yielding acetate and the hydroxylated counterparts of the original aromatics^41^ (Fig. 1A). While one of the acetate’s carbons comes from the methoxyl group, the other carbon is originated from fixed CO ^40^. In addition, *A. woodii* can also reduce any double bonds present in the possible acrylate side chains of the aromatic substrates^41,43^.

**Figure 1:**
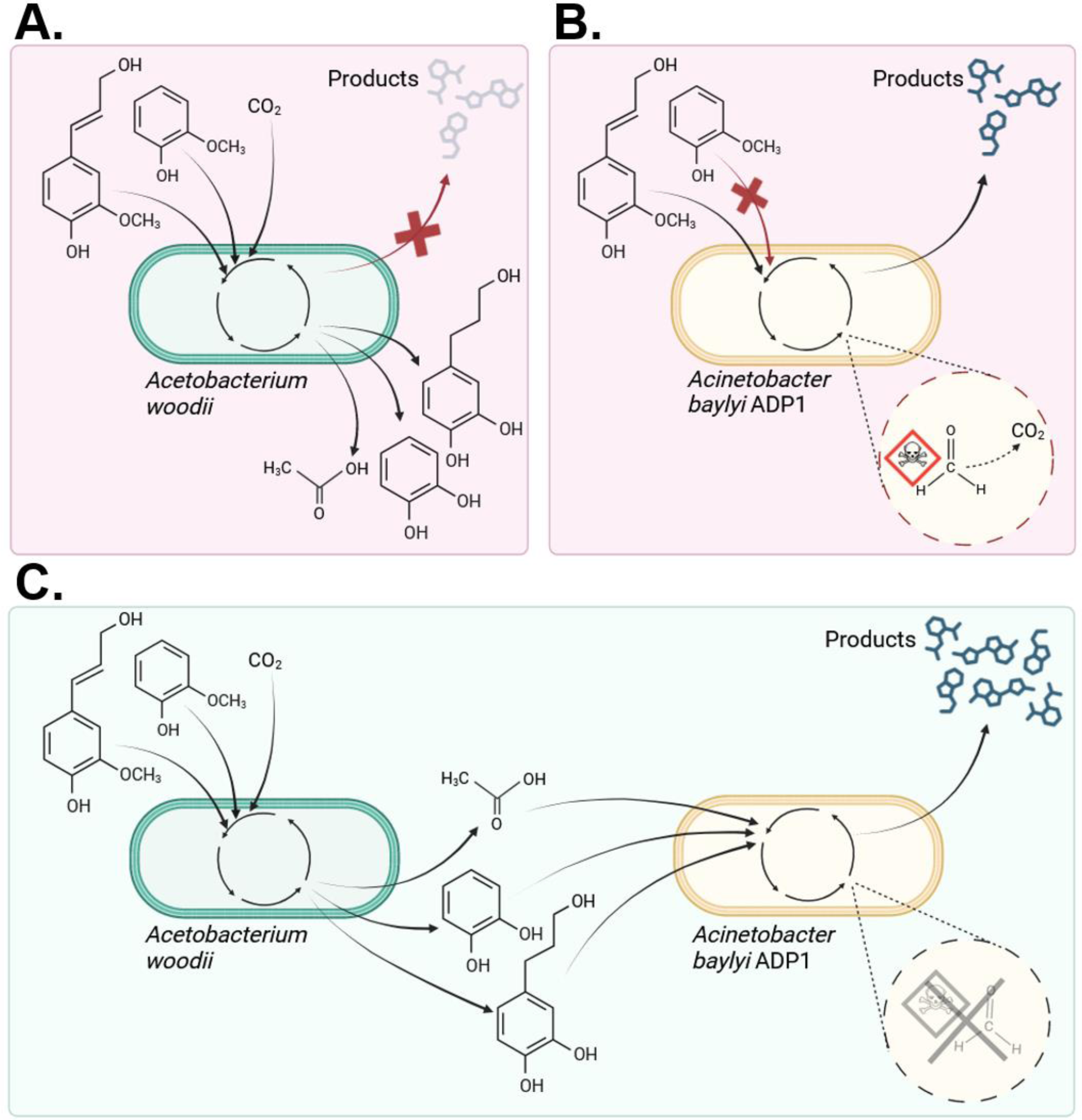
The valorization of lignin-related compounds to products by A. woodii and/or ADP1. Ferulic acid and guaiacol are used as example model compounds. A.) A. woodii utilizes methyl groups of the compounds as carbon and electron donors and double bond of the ferulic acid as electron acceptor in a process that fixates CO2 and the methyl group into acetate. However, the modified aromatic compounds cannot be further utilized by A. woodii. B.) ADP1 cannot use guaiacol for its growth, so it is left unutilized. Ferulic acid is utilized as a substrate via β-ketoadipate pathway and can be directed to biomass or products of interest, but the demethylation process produces toxic formaldehyde which is degraded into CO2 (presented with dashed circle). C.) By modifying ferulic acid and guaiacol first anaerobically with A. woodii, ADP1 can utilize all the obtained products for its substrate. Theoretically, higher yields can be obtained compared to the process including only ADP1. In addition, by executing the demethylation of the aromatic compounds anaerobically, the formation of the toxic by-product formaldehyde can be avoided.

We hypothesized that the metabolites produced by *A. woodii* when cultivated on aromatic compounds could be used as substrates by ADP1. By utilizing anaerobic demethylation, ADP1 gets an access to previously unavailable carbon: if ADP1 performed the demethylation reaction by itself, the carbon from the methoxyl group would form cytotoxic formaldehyde, eventually oxidized into CO2 (Fig. 1B). Therefore, the anaerobic demethylation increases the number of the available carbon for the cells, removes the need for the enzymatic step producing a toxic by-product, and decreases the negative effect of cofactor imbalance caused by aerobic demethylation reaction (Fig. 1C).

### The utilization of compounds modelling the anaerobic demethylation metabolites of *A. woodii* by ADP1

We first studied if ADP1 could grow on the substrates modelling the demethylation metabolites of *A. woodii*. In addition, we were interested to see if the growth metrics obtained on the metabolites would differ from the situation where ADP1 is cultivated solely on the unmodified methoxylated aromatics. According to the literature, when *A. woodii* is cultivated on 10 mM vanillate, 10 mM PCA and 7.5 mM acetate are produced^41^ (Table S1, Fig. 2A). In our experiments, molar yields of similar range were obtained (Table S2). The effects of the different carbon sources on biomass yields were first examined by performing a flux balance analysis (FBA). According to the ADP1 model, 10 mM vanillate provides 0.77 grams of cell dry weight (gCDW), whereas the biomass production from the mixture of 10 mM PCA and 7.5 mM acetate is 0.96 gCDW. The results of the FBA were supported by the experimental data: when we cultivated ADP1 on 10 mM vanillate or on 10 mM PCA and 7.5 mM acetate mixture (modelling the metabolites produced by *A. woodii* during vanillate demethylation), faster cell growth and higher OD was achieved in the latter culture (Fig. 2B). Despite acetate is known to repress the utilization of aromatic compounds in ADP1^44^, we observed no negative effects or diauxic growth pattern on the culture with a mixture of PCA and acetate.

**Figure 2:**
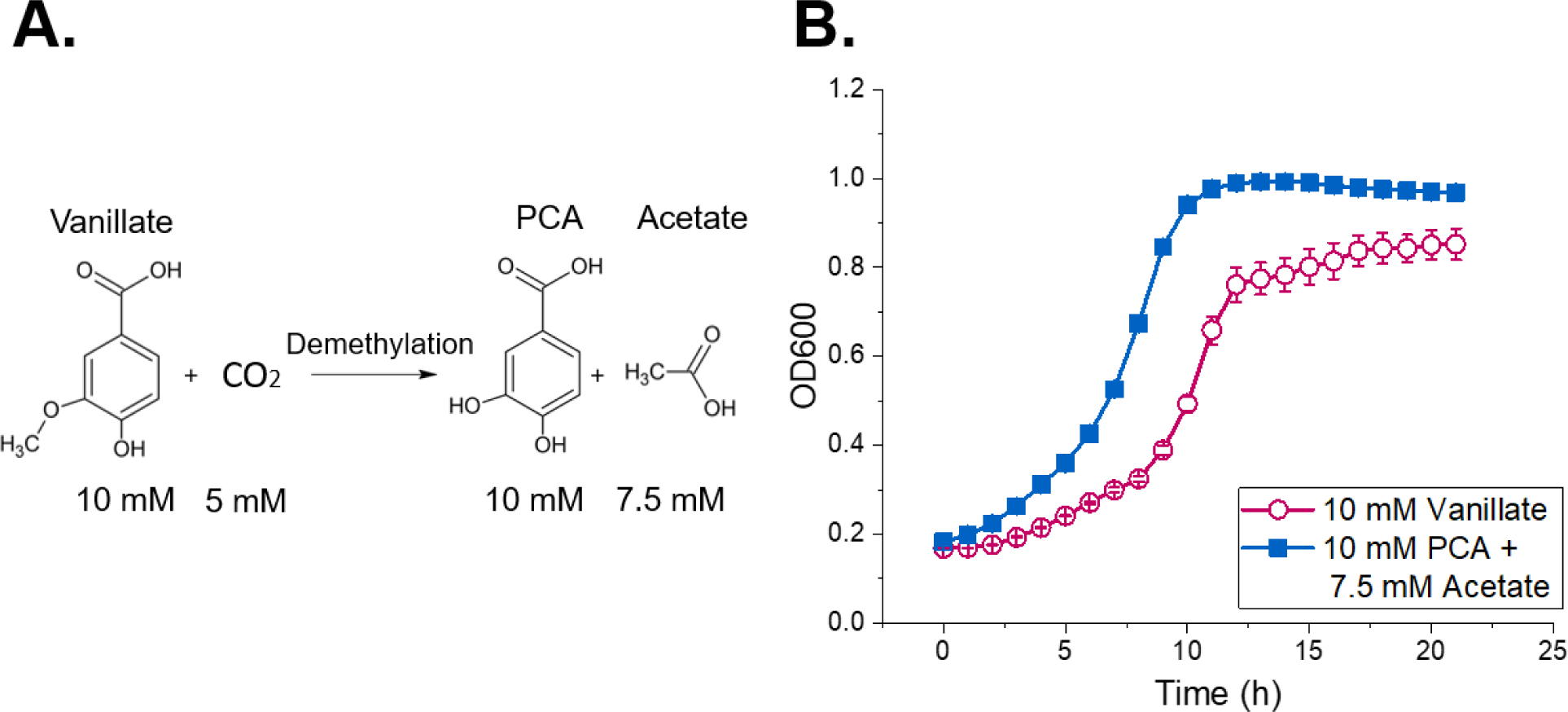
A.) Vanillate demethylation reaction in A. woodii in the presence of CO2. 10 mM vanillate is demethylated into 10 mM PCA and 7.5 mM acetate. B.) The difference of ADP1 growth on 10 mM vanillate and on mixture containing 10 mM PCA and 7.5 mM acetate. The mixture represents the end-products of vanillate demethylation by A. woodii. The error bars indicate the standard deviations from the average values of the biologically independent triplicates. In some cases, the error bars are smaller than the size of the marker.

Thus, the anaerobic conversion of the methoxylated aromatic substrates into a more accessible form, accompanied by CO2 fixation into acetate, has clear advantages, seen as both higher biomass an improved growth rate of ADP1.

### The utilization of ferulate and coumarate by *A. woodii* and ADP1

Ferulate and coumarate are model compounds of G- and H-lignin, respectively, that both contain acrylate side chains. According to the literature, *A. woodii* produces 1.0 mol dihydrocaffeate ((3-(3,4- dihydroxyphenyl)propanoate) and 0.5 mol acetate from 1.0 mol ferulate^41^. Coumarate does not contain any methoxyl groups, so the only modification is the reduction of the double bond in the acrylate tail, theoretically resulting in the formation of stoichiometric amounts of phloretate (3-(4- hydroxyphenyl)propionate)^45^. We studied the growth of *A. woodii* on 13 mM ferulate or 14 mM coumarate under N2-CO2. When ferulate was used as a substrate, we detected notable amounts of caffeate (3-(3,4-dihydroxyphenyl)-2-propenoate) in addition to dihydrocaffeate and acetate (Fig. S1A), indicating that the demethylation of the methoxyl group of ferulate along with CO2 fixation occurs preferentially over double bond reduction in the studied conditions. The molar yields of dihydrocaffeate and acetate from ferulate were 0.70 ± 0.24 and 0.66 ± 0.14, respectively (Table S3). When *A. woodii* was cultivated on coumarate, formation of phloretate (3-(4- hydroxyphenyl)propionate) and acetate was detected to some extent, despite the absence of added electron donor, indicating that energy for the double bond reduction and CO2 fixation was provided by carry-over fructose from the preculture (Fig. S1B) (Table S4).

We then investigated the growth metrics of ADP1 on ferulate and coumarate in comparison to the growth on the metabolites provided by *A. woodii* cultivated on ferulate and coumarate, namely caffeate, dihydrocaffeate, and phloretate (Fig. 3A). In this experiment, we concentrated solely on the aromatic metabolites and thus neglected the positive effect of acetate on growth (Fig. 2B). A clear difference in the growth of ADP1 was observed between the different substrates: in all cases, the obtained biomass and growth rate on demethylated and/or reduced aromatics was higher compared to those on ferulate and coumarate (Fig. 3B, 3C), and the effect was even more pronounced with higher concentrations (Fig. S2 and Fig. S3); when ADP1 was cultivated on 15 mM ferulate, the lag-phase lasted approximately one day and the obtained biomass was modest, whereas with caffeate, the exponential growth phase was reached earlier, and the biomass growth resulted in OD 1.2. The final OD of the cells grown on dihydrocaffeate was the same than with caffeate-grown cells, but the growth occurred more slowly. The difference in the cell growth on coumarate and phloretate was less drastic, although phloretate-grown cells had shorter lag-phase with final OD of 1.2, whereas with coumarate-grown cell, the final OD remained under 1.

**Figure 3:**
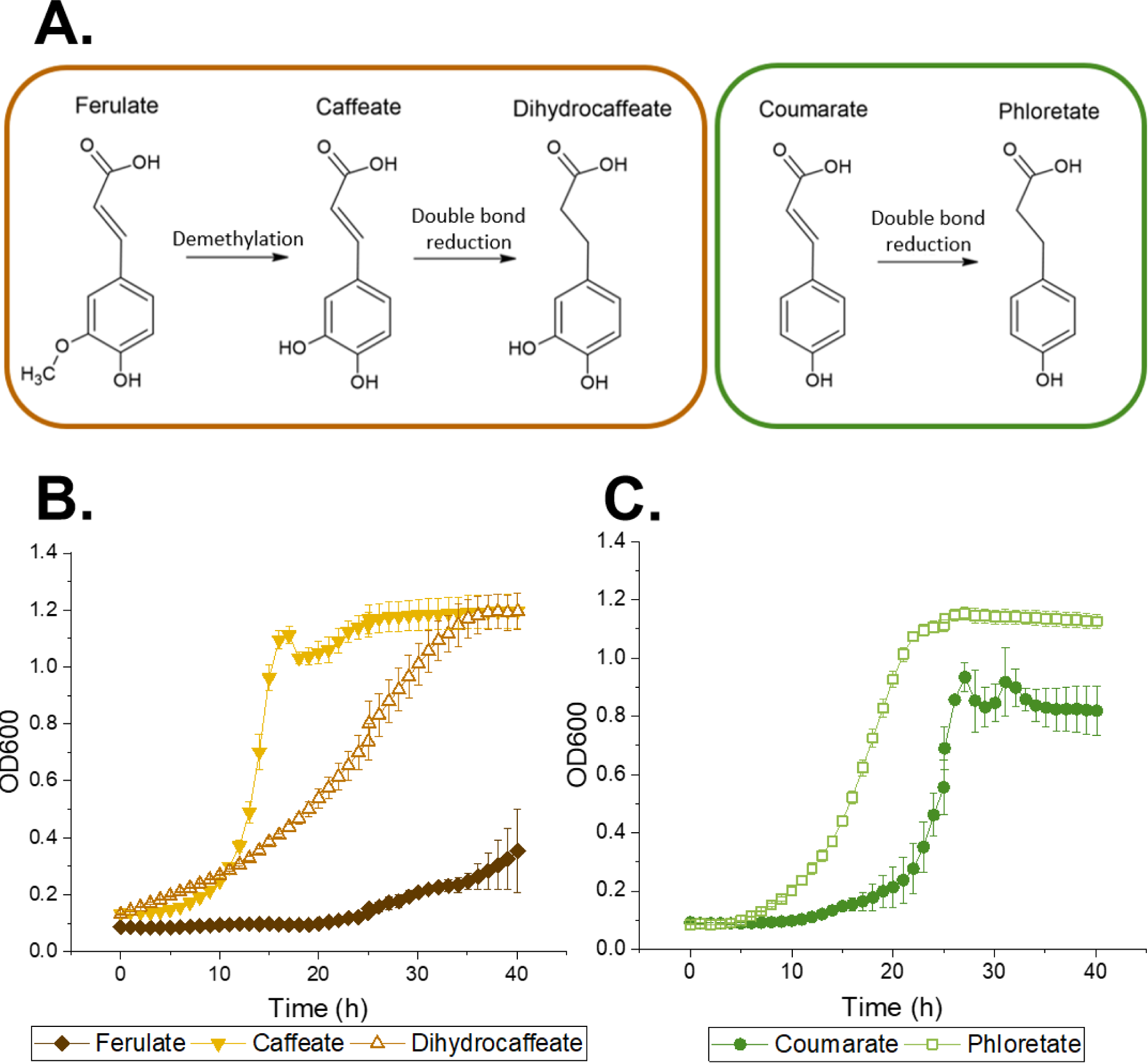
A) A simplified scheme of the modification carried out by A. woodii for ferulate and coumarate. B). The growth of ADP1 on 15 mM of ferulate, caffeate and dihydrocaffeate. C.) The growth of ADP1 on 15 mM coumarate and phloretate. The error bars in graph B and C indicate the standard deviations from the average values of the biologically independent triplicates. In some cases, the error bars are smaller than the size of the marker.

These results affirm that the modifications carried out by *A. woodii* for the aromatic compounds are beneficial for ADP1 growth: the growth metrics were enhanced, indicating that caffeate, dihydrocaffeate, and phloretate are more readily consumed and potentially less toxic carbon sources compared to ferulate and coumarate, respectively. However, interestingly, the ADP1 growth was faster on double bond-containing caffeate than on dihydrocaffeate lacking the double bond. We hypothesize the phenomenon to be related to the catabolic route of dihydrocaffeate in ADP1; unlike with caffeate, the catabolism of dihydrocaffeate requires an additional enzymatic step, where 3,4- dehydroxyphenolpropionyl-CoA is converted to caffeoyl-CoA, which is a direct degradation intermediate of caffeate catabolism^46^. This finding creates an interesting opportunity: when *A. woodii* was cultivated on ferulate, complete conversion of ferulate into dihydrocaffeate was not observed, but notable amounts of caffeate were still present in the medium after 164 h (Fig. S1A). Thus, much shorter cultivation time of *A. woodii* would be sufficient, making the overall cultivation process faster and simultaneously providing the most suitable carbon source and maximal amount of acetate for ADP1 growth.

The negative effect of the methoxyl group on ADP1 growth is evident when ferulate and caffeate are compared, because the only difference between the two compounds is the lack of the methoxyl group in caffeate^46^. Most likely the lack of methoxyl group on both coumarate and phloretate molecules explain why the growth metrics of ADP1 on these compounds did not differ that dramatically. To the best of our knowledge, there are no previous studies reporting the growth of ADP1 on phloretate. Therefore, the exact metabolic pathway for phloretate catabolism is not yet known and the reason why ADP1 grew better on phloretate than coumarate remains unclear.

### Complete utilization of guaiacol by *A. woodii* and ADP1 coculture

After confirming the suitability of *A. woodii* metabolites on ADP1 growth, we next established a two-stage culture for the complete utilization of a lignin-related model compound, guaiacol. Unlike vanillate, ferulate, or coumarate, ADP1 cannot naturally utilize guaiacol. We further confirmed this by cultivating ADP1 in MSM with guaiacol as a sole carbon source (data not shown). It has been previously reported that *A. woodii* can grow on guaiacol, producing catechol and acetate as demethylation products^47,48^. To obtain quantitative information about the process, we cultivated *A. woodii* on 6, 11, and 18 mM guaiacol as a sole organic carbon source (Table S5). The obtained OD values and the molar yields of catechol and acetate from guaiacol were in line with previous literature, being approximately 0.85 mol/mol for catechol and 0.7 mol/mol for acetate.

While ADP1 cannot grow on guaiacol, both catechol and acetate serve as suitable carbon sources for its growth. However, we were concerned about the toxicity of catechol to ADP1. For example, Kohlstedt *et al.*^49^ have reported that catechol concentrations above 2 mM caused reduced growth rates while 8 mM catechol completely inhibited the cell growth of *P. putida*. Thus, the tolerance of ADP1 towards catechol was first tested by cultivations in MSM supplemented with 0.5, 1, 2, 5, 7.5, and 10 mM of catechol as a sole carbon source (Supplementary FigS4). ADP1 showed unexpectedly high tolerance towards catechol, as the highest ODs were achieved with 5 mM and 7.5 mM catechol concentrations, which were 0.7 and 0.8, respectively. However, 7.5 mM catechol caused a notable lag-phase, and the exponential cell growth started only after 30 h of cultivation.

As no drastic differences in the obtained biomass between 5 mM and 7.5 mM catechol concentrations were detected, we decided to test the two-stage coculture of *A. woodii* and ADP1 using 6 mM guaiacol as a sole organic carbon source (Fig. 4). *A. woodii* consumed all guaiacol within 48 hours of cultivation and converted it to catechol and acetate. After the anaerobic phase, the cultivation tubes were opened and ADP1 cells were added to the culture. At the end of the aerobic cultivation with ADP1, no acetate or catechol were detected in the medium.

**Figure 4:**
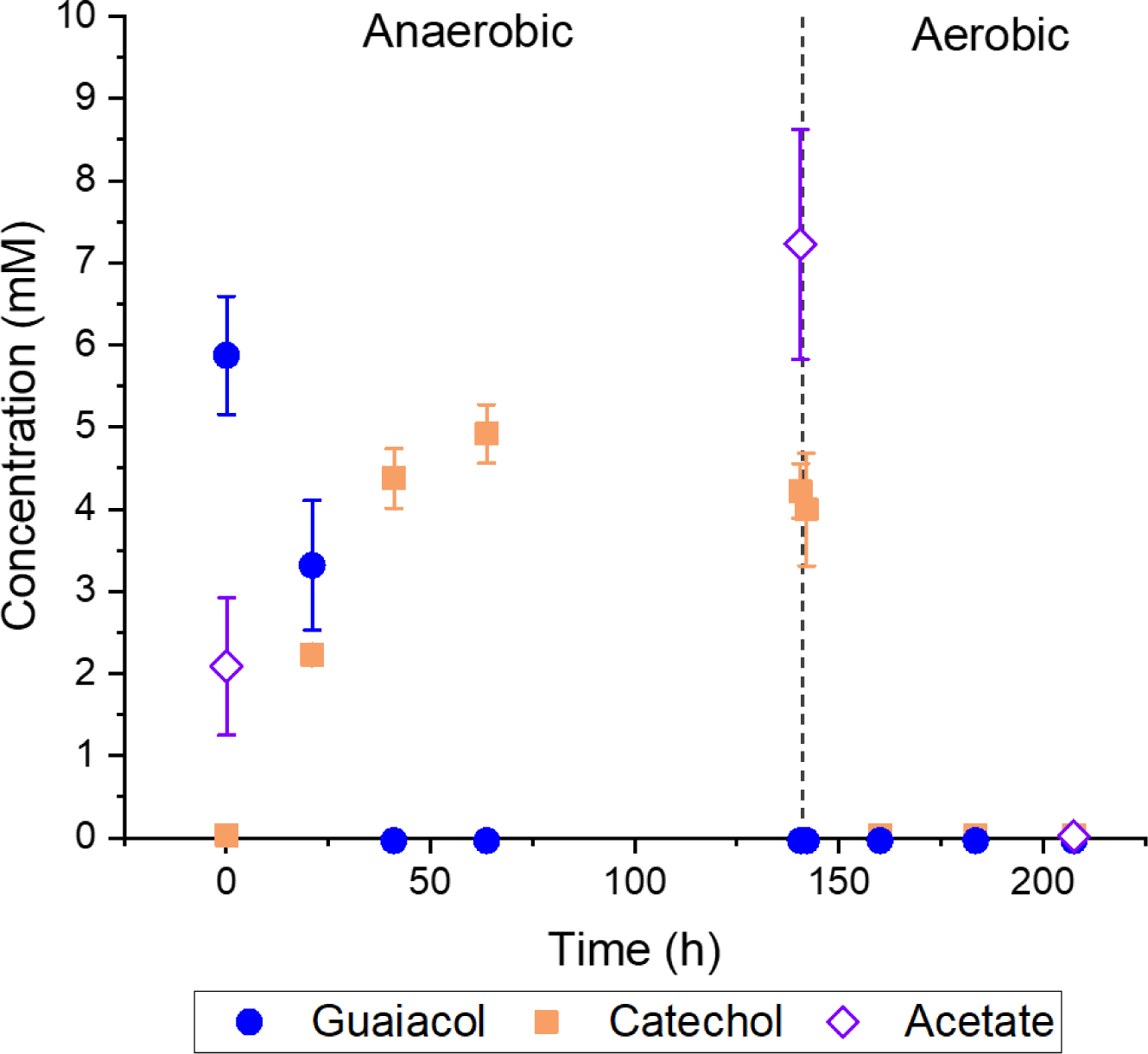
Two-stage cocultivation of A. woodii and ADP1 on acetobacterium medium supplemented with 6 mM guaiacol as the sole organic carbon source. The increase of acetate concentration at the beginning of the cultivation is a result of the carry-over acetate from A. woodii inoculation. ADP1 was inoculated later, at 141 h, marked in the graph with dashed vertical line. The error bars indicate the standard deviations from the average values of the biologically independent duplicates. In some cases, the error bars are smaller than the size of the marker.

It was confirmed that ADP1 was able to grow on the authentic metabolites produced by *A. woodii* in the acetobacterium medium (ABM). By utilizing the coculture approach, a complete degradation of guaiacol was achieved using wild-type cells, without the need for genetic engineering.

### Production of muconic acid from guaiacol with *A. woodii* and ADP1 coculture

To demonstrate that lignin-related aromatic compounds could be converted into products by utilizing the integrated metabolisms of *A. woodii* and ADP1, we established the production of MA from guaiacol. Muconic acid is a natural intermediate compound of the β-ketoadipate pathway, located in the catechol branch. To allow the accumulation of MA, we constructed a knock-out mutant strain with *catBC* deletion, thus preventing further catabolism of MA. The resulting knock-out strain was designated as ADP1ΔcatBC.

We tested the production of MA in a two-stage process, where we first cultivated *A. woodii* on 11 mM guaiacol as a sole organic carbon source. After the conversion of guaiacol to 10.9 ± 0.3 mM of catechol and 8.4 ± 1.8 mM of acetate, the anaerobic cultivation was stopped and *A. woodii* cells were removed from the cultivation by centrifugation. The cultivation medium was diluted to lower the catechol concentration to 5.4 ± 0.2 mM and ADP1ΔcatBC cells were added to the culture.

ADP1ΔcatBC was able to grow and produce MA solely from *A. woodii* metabolites. Because of the knock-out mutation, only acetate was used to produce biomass reaching the OD 1.0 ± 0.0. The strain converted all catechol into ccMA, but because of the spontaneous isomerization of the compound, also small amounts of *cis-trans* muconic acid (ctMA) was detected from the medium. We further confirmed the product to be MA with LC-MS analysis. In the diluted cultivation, the total concentration of produced MA was 4.9 ± 0.1 mM. Therefore, the total molar yield of MA from catechol was 0.91 ± 0.05 mol/mol, while the total molar yield of MA from guaiacol was 0.87 ± 0.04 mol/mol.

Previously, production of MA from guaiacol has been achieved by engineered *Amycolatopsis* sp. ATCC 39116^50^ and *P. putida*^30^ with high molar yields. However, in addition to guaiacol, the cultivations were supplemented with glucose to support primary metabolism and cell growth; Almqvist *et al.*^30^ reported that guaiacol demethylation caused a severe redox equivalent imbalance to the cells, and by adding glucose to the cultivations, 10-folds higher MA production rates were achieved. They tested whether vanillin, present in their depolymerized lignin feedstock, could replace glucose as an additional carbon source, but too low ratio of vanillin to guaiacol in the feedstock as well as inhibitory effect caused by higher vanillin concentrations were found to be problematic. In contrast, by utilizing our coculture approach, guaiacol can serve as the sole organic carbon source, as the anaerobic demethylation of guaiacol along with CO2 fixation provides acetate for ADP1 growth and circumvents the redox imbalance issue.

### A one-pot cocultivation of *A. woodii* and ADP1 for muconic acid production

While *A. woodii* and ADP1 represent strictly anaerobic and aerobic bacteria, respectively, our previous work indicates that it might be possible to carry out a one-pot cocultivation of *A. woodii* and ADP1 with alternating anaerobic and aerobic phases^51,52^. The one-pot approach allows for more straight- forward culture operation and potentially reduces the conversion time of substrates to products. To that end, we established a three-step batch process in a bioreactor for the production of MA from guaiacol.

We started the cultivation by adding ADP1ΔcatBC in ABM medium prepared without using any anaerobic techniques. In addition to 5 mM guaiacol, the medium was supplemented with 10 mM glucose to ensure the oxidation activity of ADP1 during the first phase of the cultivation, referred to as ‘deoxygenation phase’. However, the addition of glucose turned out to be unnecessary as ADP1 did not utilize glucose during the first phase (Fig. S5). After the cell inoculation, the reactor was closed, and ADP1ΔcatBC was able to rapidly deoxygenize the medium. Thereafter, *A. woodii* cells were anaerobically inoculated into the reactor, which started the anaerobic phase. After three days, *A. woodii* had completely demethylated guaiacol into catechol. Minor amounts of glucose (1.3 mM) were consumed during the anaerobic phase (Fig. S5). The final aerobic phase was initiated by starting the airflow, enabling the growth of ADP1ΔcatBC cells. ADP1ΔcatBC was able to convert all the available catechol into ccMA. During the aerobic phase, all acetate produced by *A. woodii* was completely consumed. ADP1 started to utilize glucose only after other carbon sources were depleted (Fig. S5), indicating that glucose supplementation is not needed in the process.

In the one-pot approach, the metabolic integration between the two bacteria is even more stringent compared to two-stage cultivations as both bacteria are dependent on the metabolic activities of the other: by the deoxygenation of the medium, ADP1 modified the growth environment to be suitable for *A. woodii* growth. The growth of ADP1 also produces CO2, required for the demethylation reactions of the acetogen, while ADP1 is dependent on the end-metabolites of *A. woodii* (Fig. 5A). Encouragingly, our results suggest that both bacteria were able to stay active in the bioreactor conditions: The consumption of acetate (Fig. S5) and catechol (Fig. 5C) that occurred already during the anaerobic phase indicates that ADP1ΔcatBC was metabolically active by consuming the minor amounts of oxygen that potentially leaked into the bioreactor, while keeping the medium anaerobic and suitable for *A. woodii* to grow. These exciting results suggest that indeed, it is possible to cocultivate the strictly anaerobic acetogen *A. woodii* together with a strictly aerobic bacteria, opening novel and intriguing possibilities for the utilization of *A. woodii*. To the best of our knowledge, our study represents the first example of utilizing *A. woodii* in a one-pot cocultivation containing both aerobic and anaerobic cultivation steps.

**Figure 5:**
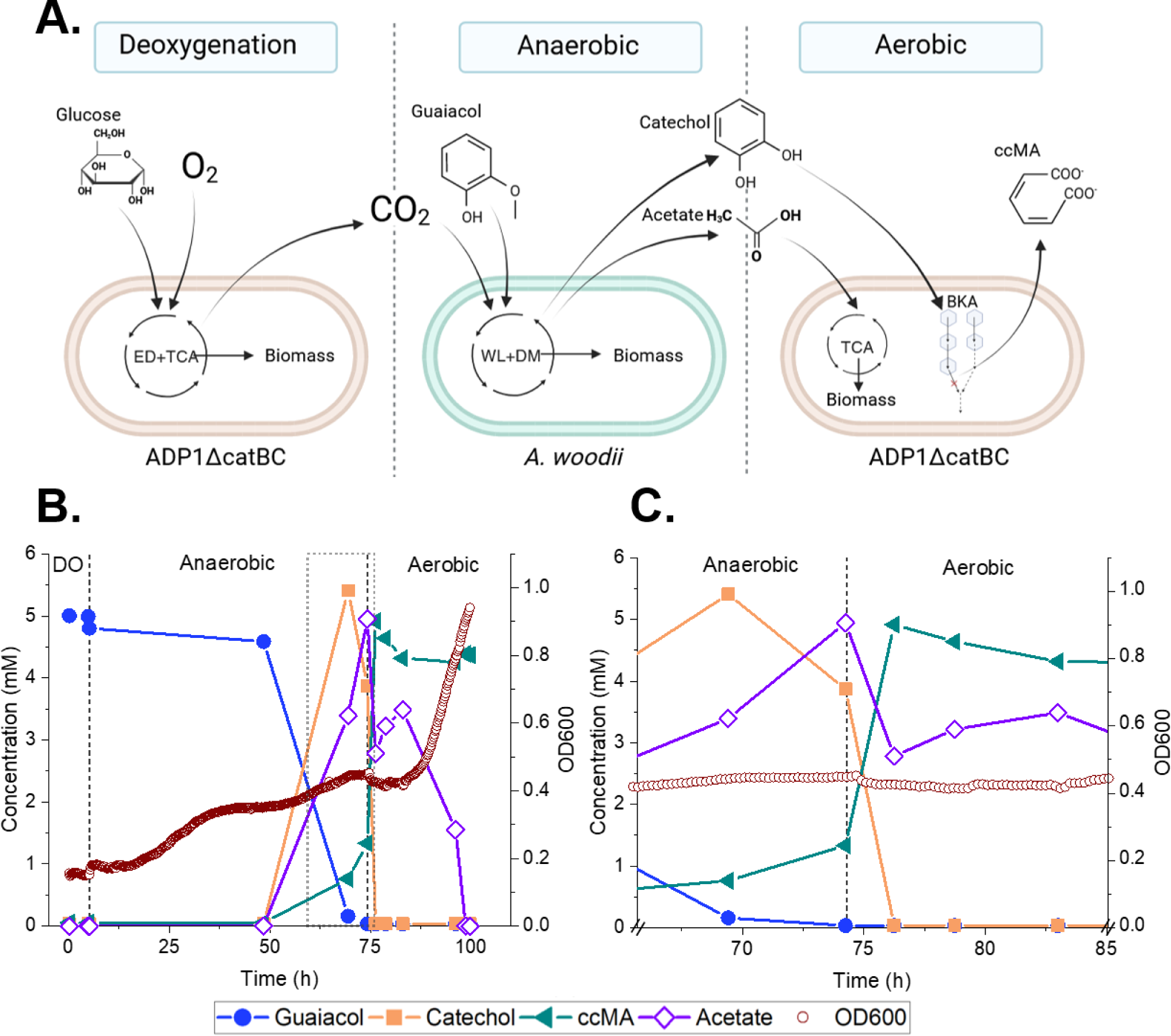
Production of MA by ADP1ΔcatBC and A. woodii one-pot cocultivation. A.) Hypothetical schematics of the cultivation phases and the substrates and products in each phase. ED: Entner-Doudoroff pathway, TCA: Tricarboxylic acid cycle, WL: Wood-Ljungdahl pathway, DM: demethylation, BKA: β-ketoadipate pathway. B.) Experimental data of the three-phase one-pot coculture of A. woodii and ADP1ΔcatBC. The carry-over acetate resulted from the inoculation of A. woodii is subtracted from the acetate concentration, which therefore describes solely the acetate produced by A. woodii during the coculture. C). Close-up of the experimental data from 65.5 h - 85.5 h. Because of the spontaneous isomerization of ccMA to ctMA, the concentration of ccMA decreases during the aerobic phase in the graphs B and C.

Equimolar conversion of guaiacol into MA in the one-pot cultivation was achieved. Notably, ADP1ΔcatBC converted catechol very fast, within few hours, into MA. The conversion of catechol to MA occurred before all the acetate was consumed (Fig. 5), indicating that the conversion is not under carbon catabolite repression (CCR) caused by acetate, unlike previously suggested^53^. In addition, ADP1 exhibited good tolerance and metabolic activity towards catechol, as no inhibition of growth or MA production was detected. These observations well justify the selection of ADP1 as the optimal strain for the coculture set-up, as catechol has been identified as the most toxic lignin-related compound to for example *P. putida,* which has been previously broadly employed for MA production^11^.

Nowadays, many of the state-of-the-art approaches to MA production rely on the heterogeneous expression of PCA decarboxylases converting PCA to catechol^54–56^. Using PCA decarboxylases such as AroY^57^ can broaden the range of suitable substrates for MA accumulation, including vanillate, ferulate, and coumarate. However, producing PCA from lignin-related methoxylated aromatics still requires the problematic step of aerobic demethylation. Thus, expressing AroY in ADP1 could be a potential approach, allowing for the production of MA also from methoxyl-free and readily accessible caffeate, dihydrocaffeate, and phloretate produced by *A. woodii*.

## Discussion

Aerobic demethylation is a crucial step in the catabolism of the methoxyl group-rich G- and S-lignin derived compounds, but the process produces cytotoxic formaldehyde causing cofactor imbalance and results in wasting the carbons from the methoxyl groups into CO2. Here, we presented a novel approach based on the optimal integration of anaerobic and aerobic metabolisms to circumvent these problems while achieving more efficient utilization of lignin-related compounds: by employing anaerobic demethylation of the acetogen *A. woodii*, the carbons from the methoxyl groups and CO2 are reduced to acetate, which together with the demethylated aromatic compounds can be utilized and upgraded to products by the aerobic bacterium *A. baylyi* ADP1. Thus, our approach converts the methoxyl group of the aromatic compound from a noxious vice into a valuable asset and makes the previously unavailable carbon accessible for much wider range of microbes while substantially enhancing the carbon recovery. Importantly, not only does the acetate contribute to the growth of aerobic ADP1, but the aromatic compounds modified by *A. woodii* are also more accessible and better tolerated by ADP1.

By employing the coculture approach, we were able to achieve utilization of guaiacol for production, which has traditionally been considered as a difficult task. By the stringent integration of the metabolisms, the consortium was able to accomplish tasks neither bacterium could perform alone: guaiacol cannot be utilized by ADP1, and *A. woodii* cannot perform ring-fission reaction required for MA production. We further demonstrated a successful cocultivation of strictly aerobic and anaerobic bacteria by establishing a simple one-pot process of *A. woodii* and ADP1ΔcatBC, by which we achieved the production of MA from guaiacol with an equimolar yield. The process required minimal genetic engineering, namely a single knock-out, to allow the accumulation of MA by ADP1ΔcatBC, thus allowing for robust and stable production system. ADP1 also showed good tolerance and high catabolic activity towards catechol, a key intermediate in the system. In addition, the acetogenic demethylation of guaiacol coupled with CO2 fixation did not only provide precursor for the product, but also the growth substrate for ADP1, eliminating the need for using additional carbon sources. Such comprehensive utilization of carbon would not be possible using only aerobic production system.

This study highlights and repurposes unique anaerobic characteristics and metabolic features that have been previously overlooked in the context of lignin valorization. By combining these features with aerobic metabolism in rationally designed synthetic cocultures, many challenges related to upgrading the recalcitrant feedstock can be tackled. Moreover, the concept of synergistically integrating metabolisms could be potentially applied across various microbial production platforms and to substrates beyond those derived from lignin.

## Materials and methods

### Strains and cultivations

*Acinetobacter baylyi* ADP1 (DSM 24193), *A. baylyi* ADP1ΔcatBC::tdk/kan, and *Acetobacterium woodii* (DSM 1030) were used in this study. ADP1 DSM24193 was used as a background strain for the knock- out strain with *CatB* (ACIAD1446) and *CatC* (ACIAD1447) deletions, referred here after as ADP1ΔcatBC. The molecular work for constructing the knock-out strain was done by using established methods. All the reagents and the primers were purchased from ThermoFisher Scientific (USA).

The three gene components for the knock-out cassette were first amplified separately by PCR as described previously^58^ with following exceptions: the upstream flanking of the cassette was constructed by amplifying the region between CatB_P3 and CatB_P4 primers from the ADP1 genome. The downstream flanking was amplified from ADP1 genome by using the primers CatC_P5 and CatC_P6. The tdk/kan^r^ ^59^ segment of the cassette, containing kanamycin resistance gene, was amplified from the genome of *Acinetobacter baylyi* ADP1Δ*acr1*::tdk/kan^r^ ^60^ (a kind gift of Veronique de Berardinis) by using the primers Tdk_kanF and Tdk_kanR. The knock-out cassette was constructed by using overlap extension PCR, and the complete cassette was amplified by using primers CatB_P3 and CatC_P6. The sequence of the obtained cassette and the used primers are presented in the supplementary material. The knock-out cassette was transformed to ADP1 as described previously^61^. The kanamycin concentration was 25 µg/ml.

Unless stated otherwise, ADP1 was precultivated in 5 ml of minimal salt medium (MSM)^62^ containing K2HPO4 (3.88 g/L), NaH2PO4 (1.63 g/L), (NH4)2SO4 (2.00 g/L), MgCl2 * 6 H2O (0.1 g/L), EDTA (10 mg/L), ZnSO4 * 7H2O (2 mg/L), CaCl2 * 2 H2O (1 mg/L), FeSO4 * 7 H2O (5 mg/L), Na2MoO4 * 2 H2O (0.2 mg/L), CuSO4 * 5 H2O (0.2 mg/L), CoCl2 * 6 H2O (0.4 mg/L) and MnCl2 * 2 H2O (1 mg/L), supplemented with 0.2% casamino acids and 0.4% D-glucose. For ADP1ΔcatBC precultivations, also 25 µg/ml kanamycin was added. ADP1 and ADP1ΔcatBC cultivations were done at +30⁰C and 300 rpm, unless stated otherwise. 96-well plate cultivations were done by using Tecan Spark multimode microplate reader (Tecan, Switzerland) with 96-well µClear®plates (Greiner bio-one, Germany). The cultivation volume was 200 µl and the temperature was kept at +30⁰C. Shaking amplitude was 2.5 mm with the frequency of 108 rpm, and the shaking duration was 180 seconds. Shaking was performed before and after the measurements of optical density at 600 nm, which occurred once in an hour.

Unless stated otherwise, *A. woodii* was cultivated in 12 ml of acetobacterium medium (DSMZ 135, referred as ABM) which was prepared according to DSMZ’s instructions. The medium was prepared under N2-CO2 80:20 headspace and the final composition of the medium was the following (g/L): NaHCO3 (11.04), D-Fructose (9.79), yeast extract (1.96), NH4Cl (0.98 g/L), L-Cysteine HCl x H2O (0.49), Na2S x 9 H2O (0.49), K2HPO4 (0.44), KH2PO4 (0.32), MgSO4 x 7 H2O (0.06), nitrilotriacetic acid (0.03), NaCl (0.02), MnSO4 x H2O (0.01), CoSO4 x 7 H2O (0.004), ZnSO4 x 7 H2O (0.004), CaCl2 x 2 H2O (0.002), FeSO4 x 7 H2O (0.002), NiCl2 x 6 H2O (5.9e-4), Na-resazurin, (4.9e-4), AlK(SO4)2 x 12 H2O (3.9e-4), H3BO3 (2.0e-4), Na2MoO4 x 2 H2O (2.0e-4), CuSO4 x 5 H2O (2.0e-4), pyridoxine hydrochloride (9.8e-5), nicotinic acid (4.9e-5), p-aminobenzoic acid (4.9e-5), (DL)-alpha-lipoic acid (4.9e-5), calcium D-(+)-pantothenate (4.9e-5), thiamine HCl (4.9e-5), riboflavin (4.9e-5), biotin (2.0e-5), folic acid (2.0e-5), Na2WO4 x 2 H2O (7.8e-6), Na2SeO3 x 5 H2O (5.9e-6) and vitamin B12 (9.8e-7). Unless stated otherwise, *A. woodii* cultivations were performed at +30⁰C and 150 rpm. The inoculations for every cultivation containing *A. woodii* cells were done from fructose-grown cultivations, which were less than one week old. The inoculation volume was 0.2 ml, unless stated otherwise.

### Flux balance analysis

FBA was done by using COBRApy Python package (version 0.28.0)^63^. The metabolic network model for ADP1 was obtained from Durot *et al.*^64^ and it was used to examine the effects of using the products from anaerobic demethylation process as substrates. To perform that, new exchange reactions related to PCA intake were added to the model.

### Cultivations

The growth test of ADP1 on 10 mM vanillate and on a mixture containing 10 mM PCA and 7.5 mM acetate as well as the test where ADP1 was cultivated on 0, 5, 7.5, 10 and 15 mM of ferulate, caffeate, dihydrocaffeate, coumarate or phloretate were done on MSM in 96-well plate setting. The starting OD was set to be 0.1.

The growth of *A. woodii* on vanillate, ferulate, coumarate and guaiacol was studied on ABM without fructose supplementation. For vanillate and guaiacol, the tested concentrations were 6, 12 and 18 mM. For ferulate and coumarate, the concentrations were 13 mM and 14 mM, respectively.

The utilization of guaiacol by ADP1 was studied with 96-well plate setting by cultivating ADP1 on 0, 2.5 and 5 mM of guaiacol in MSM. Also, the growth of ADP1 on 0, 0.5, 1, 2, 5, 7.5 and 10 mM catechol was studied in similar cultivation. In both studies, the starting OD was set to be 0.1.

The ability of ADP1 to grow on the catechol and acetate produced by *A. woodii* from guaiacol was determined in a cultivation study, where *A. woodii* was cultivated first on 12 mM guaiacol in a 12.5 ml volume without fructose supplementation. After all the guaiacol was demethylated into catechol, the cultivations were transformed into sterile 50 ml Falcon tubes with vented lids. After the pH was adjusted to 7 by using 5% HCl, ADP1 cells were added to the cultivation at OD600 0.1. The pH was adjusted again to 7 on the second day of ADP1 cultivation phase.

The production of ccMA with ADP1ΔcatBC and *A. woodii* cocultivation was tested by first cultivating *A. woodii* on 11 mM guaiacol in ABM without fructose supplementation. After the anaerobic cultivation phase, the *A. woodii* cells were removed from the medium by centrifugation. To ensure that the catechol concentration would not be too high for ADP1ΔcatBC to grow and to prevent pH increase in aerobic phase caused by the presence of NaHCO3, we diluted the medium 1:1 with MQ and added MOPS buffer to a final concentration of 120 mM. In addition, 0.2% casamino acids were added to the cultivations to ensure that ADP1 would not suffer from the presence of L-cysteine^35^. The final volume of the new medium was 15 ml. ADP1ΔcatBC cells were collected from liquid preculture by centrifugation, resuspended into MSM and added to the medium to obtain a starting OD 0.3.

The three-step batch cocultivation was done by using Biostat B Plus bioreactor (Sartorius, Germany) equipped with pH-meter (Hamilton, Switzerland) and pO2-meter (Hamilton, Switzerland). Aerobically prepared low-NaHCO3 ABM (NaHCO3 concentration only 1.25 g/L) without fructose supplementation was used as a medium. The only carbon sources and electron donors added to the medium in the beginning were 10 mM glucose and 5 mM guaiacol. Before the addition of ADP1ΔcatBC cells, the medium was aired with pressured air for couple of hours. To obtain a required volume of ADP1ΔcatBC to reach the desired starting OD of 0.3, ADP1ΔcatBC was cultivated on similarly to the normal preculture, but without kanamycin and the cultivation volume was increased to 50 ml in flask. The cells were centrifugated and resuspended into 10 ml MSM. The pO2-value was calibrated to 100% just before the resuspended ADP1ΔcatBC cells were added to the reactor. The flow of the pressured air was stopped, and the reactor was sealed with clamps. The reactor was covered to protect the light- sensitive guaiacol. When pO2-value was 0.7%, *A. woodii* cells were added to the reactor, which initiated the anaerobic phase. HPLC results were used to determine the optimal starting time of the last aerobic phase. Throughout the cultivation, OD was measured in every 12 minutes by Arc view device (Hamilton, USA) and manually from each sample taken. To minimize the effect of the resazurin indicator to the OD results, manually measured OD values were used as a starting point and a gradual step-by-step transformation from one value to the next one was done based on the tendencies of machine measured values. The pH was set to 7.0 for the whole cultivation and the required adjustments were done automatically by using concentrated NaOH and H3PO4 stocks.

### Analytical techniques

Quantitative analysis of compounds was done using high performance liquid chromatography (HPLC) instruments (Shimadzu, Japan) equipped with PDA and RID detectors.

Aromatic compounds were analyzed on Luna C18 column (150 x 4.6 mm, 5 µm) (Phenomenex, USA) at 40⁰C using the mixture of MQ/Methanol/formic acid (80/20/0.16 v/v/v) as the eluent at the flow rate 1 ml/min. The injection volume was 5 µl. Aromatic substances were detected at the wavelength 254 nm or 272 nm.

Sugar and organic acid were analyzed on Rezex RHM-Monosaccharide H+ (8%) column (300 x 7.8 mm) (Phenomenex, USA) at 40⁰C using 5 mM H2SO4 as the eluent with the flow rate 0.5 ml/min. The injection volume was 10 µl. Compounds were detected by RID.

The obtainment of ccMA and ctMA in the two-stage MA production test with A. woodii and ADP1ΔCatBC was confirmed by liquid chromatography mass spectrometry (LC-MS) by using Agilent 1260 Infinity II (USA) which was equipped with UV detector and connected with Jeol Accu TOF LCPlus (JMS-T100LP) (Japan). Compounds were analyzed on Poroshell 120 EC-C18 column (100 x 4.6 mm, 2.7µ) (USA). The temperature of the column was 30⁰C, the injection volume was 2 µl. The mixture of 0.1% formic acid and methanol (80/20 v/v) was used as eluent with the flow rate 0.2 ml/min. Calculation of ccMA and ctMA was performed from extracted ion chromatograms (EIC) and UV absorption chromatograms at 272 nm using external standard method. In both cases concentration of MA was calculated as approximately 141 g/mol.

### Data availability

The authors declare that the data supporting the findings of this study are available within the paper and its supplementary information files. Additional data sets regarding the study are available from the corresponding authors on reasonable request.

## Acknowledgements

This work was financially supported by the Research Council of Finland (grant numbers 347204, 353587). SS and VS would also like to thank the Novo Nordisk Foundation (grants NNF21OC0067758, NNF22OC0079579). The authors are grateful to Dr. Pauli Losoi for his help with FBA of ADP1 and Dr. Milla Salmela for constructing the knock-out strain *A. baylyi* ADP1ΔcatBC. The figures 1A and 5 were created with BioRender.com.

## Author contributions

S.S., V.S., and E.M. designed the experimental set-up. E.M. performed the experimental work, interpreted the data, and wrote the original draft of the manuscript. E.E contributed to metabolite analyses and performed LC-MS analysis. S.S. supervised the study. E.M., V.S., and S.S. participated in reviewing and editing the manuscript. All authors read and approved the final version of the manuscript.

## Competing interests

Authors declare that there are no competing interests associated with the manuscript.

## Additional information

Supplementary information available

Correspondence and request for material should be addressed to Suvi Santala.

## Notes

### Competing Interest Statement

The authors have declared no competing interest.

