## Supplementary file for "Novel approach for carbon-wise utilization of lignin-related compounds by synergistically employing anaerobic and aerobic bacteria"

### Supplementary notes

**Cultivating *A. woodii* on vanillate as a sole organic carbon source**

The cultivation of *A. woodii* in acetobacterium medium on 6, 12 and 18 mM vanillate as a sole carbon source was done to determine if the experimental molar yields have notable difference with the literature values (Table S2). All vanillate was demethylated in every cultivation, and protocatechuate (PCA) and acetate were produced. The obtained maximum optical densities measured at 600 nm (hereafter ODs) were positively correlated with the initial vanillate concentrations, and no growth inhibition was observed with any concentrations. When *A. woodii* was cultivated without any added carbon sources, carry-over substrates from the cell inoculation resulted in minor cell growth (OD < 0.1).

**Cultivating *A. woodii* on ferulate and coumarate as a sole organic carbon source**

To study what metabolites and yields could be obtain by cultivating *A. woodii* on ferulate or coumarate and how the obtained results compare to the existing literature, *A. woodii* was cultivated on 13 mM ferulate or in 14 mM coumarate in $N_2$-$CO_2$-headspace. Acetate, caffeate (3-(3,4-dihydroxyphenyl)-2-propenoic acid) and dihydrocaffeate were obtained as metabolites, when *A. woodii* was cultivated on ferulate (Fig. 3B). The molar yields of dihydrocaffeate and acetate were 0.70 ± 0.24 and 0.66 ± 0.14, respectively (Supplementary Table S3). The conversion of caffeate to dihydrocaffeate was observed to occur at notably different rates. When *A. woodii* was cultivated on 14 mM coumarate in $N_2$-$CO_2$-headspace, phloretate (3-(4-hydroxyphenyl)propionic acid) and acetate were produced (Fig3C) with molar yields of 0.20 ± 0.05 and 0.19 ± 0.04, respectively (Supplementary Table S4).

Unlike in the literature, a complete conversion of ferulate into dihydrocaffeate with a molar yield of 1.0[1] was not achieved, although all ferulate was consumed. Dihydrocaffeate can be obtained from ferulate if ferulate's methoxyl groups is demethylated into hydroxyl group and the double bond in the acrylate side chain is reduced. The conversion of ferulate into caffeate requires less modifications: the caffeate molecule still contains the double bond in the side chain, and therefore only the methoxyl group of the ferulate must be demethylated. It can be hypothesized that the lower molar yield of dihydrocaffeate obtained in this study was resulted from the incomplete conversion of caffeate into dihydrocaffeate, most likely caused by the additional electron acceptor $CO_2$[2]. The molar yield of acetate in this study being higher than the literature value 0.5 indicated that some of the available electron from the ferulate's methoxyl groups or carry-over fructose were donated to $CO_2$ fixation.

Only 2.6 mM coumarate was consumed during the 7-day cultivation period. However, theoretically, no coumarate should have been consumed, due to the lack of electron donor groups in coumarate molecule. The reduction of coumarate into phloretate and the appearance of acetate can be explained by the carry-over substrate fructose providing electrons for the double bond reduction and $CO_2$ fixation.

**Cultivating *A. woodii* on guaiacol**

To determine the possible differences between the existing literature values and experimental data, *A. woodii* was cultivated on 6, 11 or 18 mM guaiacol in acetobacterium medium (Table S5). *A. woodii* was able to grow on guaiacol as a sole carbon source and no residual guaiacol was detected in the cultivations in the end. Catechol and acetate were detected as only end-products. When *A.* woodii was cultivated without any carbon source, the carry-over substrate fructose from the inoculation resulted only small production of acetate, 0.6 ± 0.1 mM and minor increase in biomass (final OD < 0.1 ± 0.0, increase from the initial OD only 0.05 ± 0.0).

The obtained OD values and the initial guaiacol concentrations were positively correlated, except in cultivations containing 18.1 mM guaiacol, where the biomass obtainment was similar to cultivations containing 11.4 mM guaiacol. Thus, the highest guaiacol concentration 18.1 mM seemed to hinder the biomass accumulation. This effect was also reported in the previous literature[3,4]. However, the increase in guaiacol concentrations did not result in lower molar yields of catechol like reported previously[3]. The molar yield of catechol obtained with 18.1 mM guaiacol was also noticeably higher (0.85 ± 0.0) than the catechol yield reported in literature when *A. woodii* was cultivated on 15 mM guaiacol (0.52)[3]. The variances in the obtained molar yields of acetate was greater than in the molar yields of catechol, but the acetate concentrations are in the same range than reported in the literature[4].

**Sequence of the knockout cassette:**

acagtaaggagctgacgtaaccaattctcaaggtttgactgaccgttgcaatccgttttgccattgaggacgcttgtgcagtatgagtcaaaatctgc
acagcatgctgataaaaaaacatgcctgcttcagtcactttagccggtctgaagccgcgttcaaatagctgaattcccaattcttcttcgagtttttga
atttgtcggctgaggggcggctgggcaatacacaacttttcagcagctttggaaatgctttgctcttcaaccacggtcacaaaatatctgaggtgtct
tagttccatttatacgccctaattggttttatataccttttttagtatgcaaaaataccaaattgtttatctttttattattacattaatttaaggtatgtaaa
tagtatttattgaaaagaagatggaccgatgtataaatcagtggaaactattttaatttatttgctttattaaagaggagaaattaattaatggcaca
gctatatttctactattccgcaatgaatgcgggtaagtctacagcattgttgcaatcttcatacaattaccaggaacgcggcatgcgcactgtcgtat
atacggcagaaattgatgatcgctttggtgccgggaaagtcagttcgcgtataggtttgtcatcgcctgcaaaattatttaaccaaaattcatcattat
ttgatgagattcgtgcggaacatgaacagcaggcaattcattgcgtactggttgatgaatgccagtttttaaccagacaacaagtatatgaattatcg
gaggttgtcgatcaactcgatatacccgtactttgttatggtttacgtaccgattttcgaggtgaattatttattggcagccaatacttactggcatggtc
cgacaaactggttgaattaaaaaccatctgtttttgtggccgtaaagcaagcatggtgctgcgtcttgatcaagcaggcagaccttataacgaaggt
gagcaggtggtaattggtggtaatgaacgatacgtttctgtatgccgtaaacactataaagaggcgttacaagtcgactcattaacggctattcagg
aaaggcatcgccacgattaacggccgccaccgcggtggagctcggtacccggggatcctctagagcggacccgggaaagccacgttgtgtctca

aaatctctgatgttacattgcacaagataaaaatatatcatcatgaacaataaaactgtctgcttacataaacagtaatacaaggggtgttatgagccatattcaacgggaaacgtcttgctcgaggccgcgattaaattccaacatggatgctgatttatatgggtataaatgggctcgcgataatgtcgggcaatcaggtgcgacaatctatcgattgtatgggaagcccgatgcgccagagttgtttctgaaacatggcaaaggtagcgttgccaatgatgttacagatgagatggtcagactaaactggctgacggaatttatgcctcttccgaccatcaagcattttatccgtactcctgatgatgcatggttactcaccactgcgatccccgggaaaacagcattccaggtattagaagaatatcctgattcaggtgaaaatattgttgatgcgctggcagtgttcctgcgccggttgcattcgattcctgtttgtaattgtccttttaacagcgatcgcgtatttcgtctcgctcaggcgcaatcacgaatgaataacggtttggttgatgcgagtgattttgatgacgagcgtaatggctggcctgttgaacaagtctggaaagaaatgcataagcttttgccattctcaccggattcagtcgtcactcatggtgatttctcacttgataaccttatttttgacgaggggaaattaataggttgtattgatgttggacgagtcggaatcgcagaccgataccaggatcttgccatcctatggaactgcctcggtgagttttctccttcactacagaaacggctttttcaaaaatatggtattgataatcctgatatgaataaattgcagtttcatttgatgctcgatgagtttttctaagcatgcggagctggtatgtaaatagagatgacacttcatcagtgttcatctcttacgtttatggtatttctttattttccgttttcatccaaaggttgatgacatgatagataaaagtgcagcgaccctaacggaagcgctctcccagatccacgacggtgccaccatcctgattggtggttttggaacagccggccaacccgccgagctgattgacggactgattgaactaggtcgcaagaacctgaccatcgtcagcaacaacgccggcaatggagactatgg

### Supplementary tables

*Table S1: Stoichiometric equations of vanillate, syringate and ferulate utilization by A. woodii according to Bache and Pfennig*[1]

| Aromatic compound | Stoichiometric equation | Molar ratio of acetate/substrate | Molar ratio of acetate/$CH_3$-groups |
|---|---|---|---|
| **Vanillate** | $4\,vanillate + 2\,CO_2 + 2\,H_2O \rightarrow 4\,PCA + 3\,CH_3COOH$ | 0.75 | 0.75 |
| **Syringate** | $2\,syringate + 2\,CO_2 + 2\,H_2O \rightarrow 2\,gallate + 3\,CH_3COOH$ | 1.5 | 0.75 |
| **Ferulate** | $2\,ferulate + 2\,H_2O \rightarrow 2\,dihydrocaffeate + CH_3COOH$ | 0.5 | 0.5 |

*Table S2: The production of PCA and acetate by A. woodii when cultivated on vanillate as a sole organic carbon source. Standard deviations describe the differences between the duplicates.*

| Initial vanillate concentration (mM) | OD* | Produced PCA (mM) | Produced acetate (mM) | Molar yield of PCA from vanillate | Molar yield of acetate from vanillate |
|---|---|---|---|---|---|
| **5.9 ± 0.3** | 0.2 ± 0.0 | 4.7 ± 0.8 | 5.6 ± 0.5 | 0.80 ± 0.18 | 0.95 ± 0.13 |
| **12.2 ± 0.7** | 0.3 ± 0.0 | 11.7 ± 0.9 | 8.3 ± 1.4 | 0.96 ± 0.17 | 0.68 ± 0.15 |
| **18.4 ± 0.3** | 0.4 ± 0.0 | 14.5 ± 0.7 | 9.7 ± 0.9 | 0.79 ± 0.05 | 0.53 ± 0.06 |

*The presented OD value is the highest measured value

*Table S3: The production of caffeate, dihydrocaffeate and acetate by A. woodii when cultivated on ferulate as a sole organic carbon source*. Standard deviations describe the differences between the duplicates.*

| Initial ferulate concentration (mM) | Maximum OD | Produced caffeate (mM) | Produced dihydro-caffeate (mM) | Produced acetate (mM) | Molar yield of dihydrocaffeate from ferulate | Molar yield of acetate from ferulate |
|---|---|---|---|---|---|---|
| **12.8 ± 0.4** | 0.7 ± 0.1 | 5.6 ± 3.3 | 9.0 ± 2.9 | 8.4 ± 1.5 | 0.70 ± 0.24 | 0.66 ± 0.14 |

*Ferulate was a sole organic carbon source, but the inoculation of the cells also caused small amounts of carry-over fructose to be present in the medium.

*Table S4: The production of phloretate and acetate by A. woodii when cultivated on coumarate as a sole organic carbon source*. Standard deviations describe the differences between the duplicates.*

| Initial coumarate concentration (mM) | Maximum OD | Produced phloretate (mM) | Produced acetate** (mM) | Molar yield of phloretate from coumarate | Molar yield of acetate from coumarate |
|---|---|---|---|---|---|
| **14.1** | 0.4 ± 0.0 | 2.9 ± 0.7 | 2.7 ± 0.5 | 0.20 ± 0.05 | 0.19 ± 0.04 |

*Coumarate was a sole organic carbon source, but the inoculation of the cells also caused small amounts of carry-over fructose to be present in the medium.

**Because the acetate concentration decreased slightly towards the end of the cultivation, the concentration of the produced acetate was determined by taking an average of the last three concentrations and subtracting the concentration obtained from carry-over acetate coming from the inoculation.

*Table S5: The production of catechol and acetate by A. woodii when cultivated on guaiacol as a sole organic carbon source. The standard deviations describe the differences between the duplicates.*

| Initial guaiacol concentration (mM) | OD* | Produced catechol (mM) | Produced acetate (mM) | Molar yield of catechol | Molar yield of acetate |
|---|---|---|---|---|---|
| **5.9 ± 0.7** | 0.2 ± 0.0 | 4.9 ± 0.4 | 5.1 ± 2.2 | 0.83 ± 0.2 | 0.86 ± 0.5 |
| **11.4 ± 0.3** | 0.3 ± 0.1 | 9.7 ± 0.2 | 8.0 ± 1.1 | 0.85 ± 0.0 | 0.70 ± 0.1 |
| **18.1 ± 0.7** | 0.3 ± 0.0 | 15.3 ± 0.0 | 10.7 ± 0.4 | 0.85 ± 0.0 | 0.59 ± 0.0 |

*The presented OD value is the highest measured value

*Table S6: List of the used primers and their names*

| Name | Primer |
|---|---|
| **CatB_P3** | ACAGTAAGGAGCTGACGTAACC |
| **CatB_P4** | TTTTTATGATTTGAATTGGAGGCTGGGCGGTCCATCTTCTTTTCAA |
| **CatC_P5** | CGATGAGTTTTTCTAAGCATGCGGAGCTGGTATGTAAATAGAGATGACACTTC |
| **CatC_P6** | CCATAGTCTCCATTGCCGGC |
| **Tdk_kanF** | CCCAGCCTCCAATTCAAATCATAAAAAATTTATTTG |
| **Tdk_kanR** | CCAGCTCCGCATGCTTAGAAAAAC |

### Supplementary figures

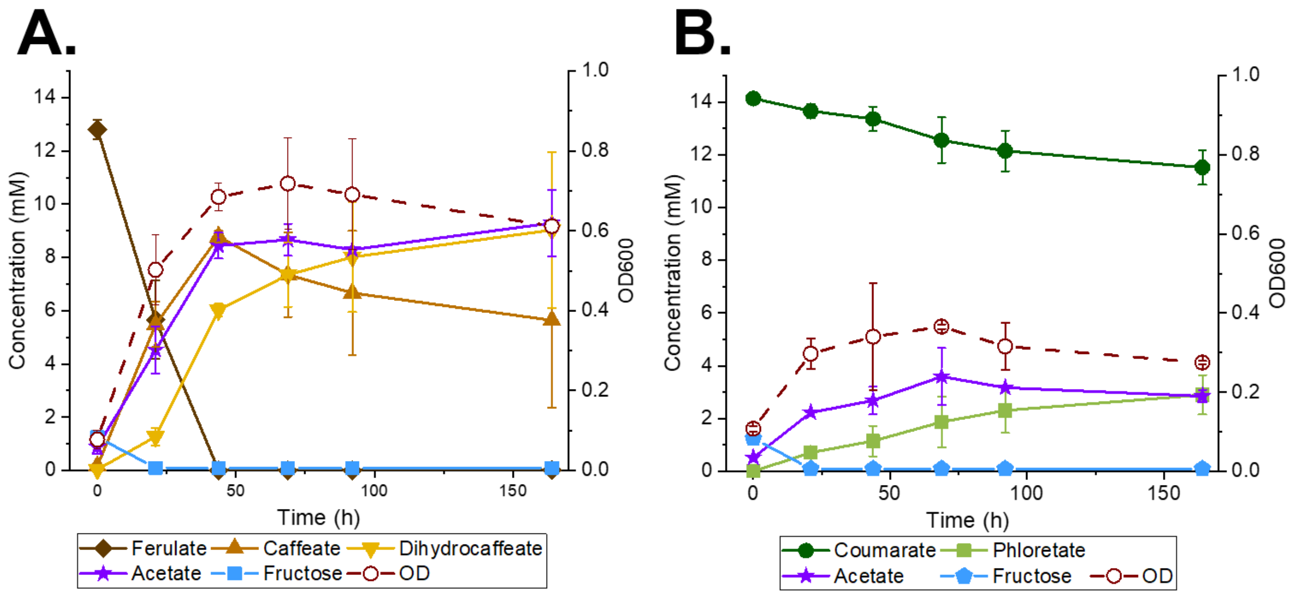


*Figure S1: A.) A. woodii cultivated in acetobacterium medium with A.) ferulate or B.) coumarate as a sole organic carbon source. The fructose present at the beginning of the cultivation is a carry-over substrate coming from the inoculation. Error bars in both graphs are describing the differences between the duplicates. The error bars indicate the standard deviations from the average values of the biologically independent duplicates. In some cases, the error bars are smaller than the size of the marker.*

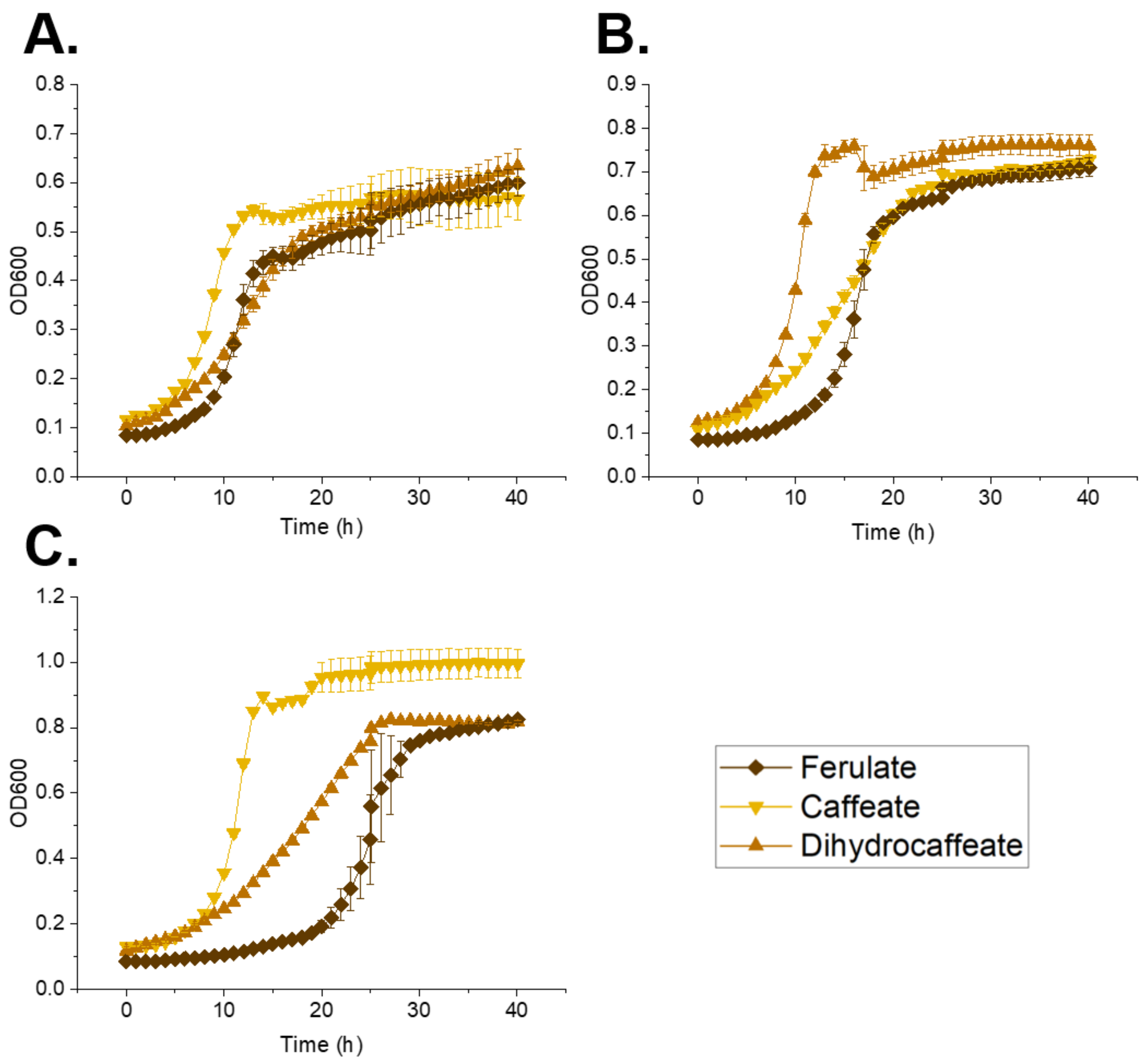


*Figure S2*: *The growth of ADP1 on ferulate or caffeate and dihydrocaffeate modelling the aromatic metabolites obtained when A. woodii is cultivated on ferulate. Used concentrations are A.) 5 mM, B.) 7.5 mM and C.) 10 mM. The error bars indicate the standard deviations from the average values of the biologically independent triplicates. In some cases, the error bars are smaller than the size of the marker.*

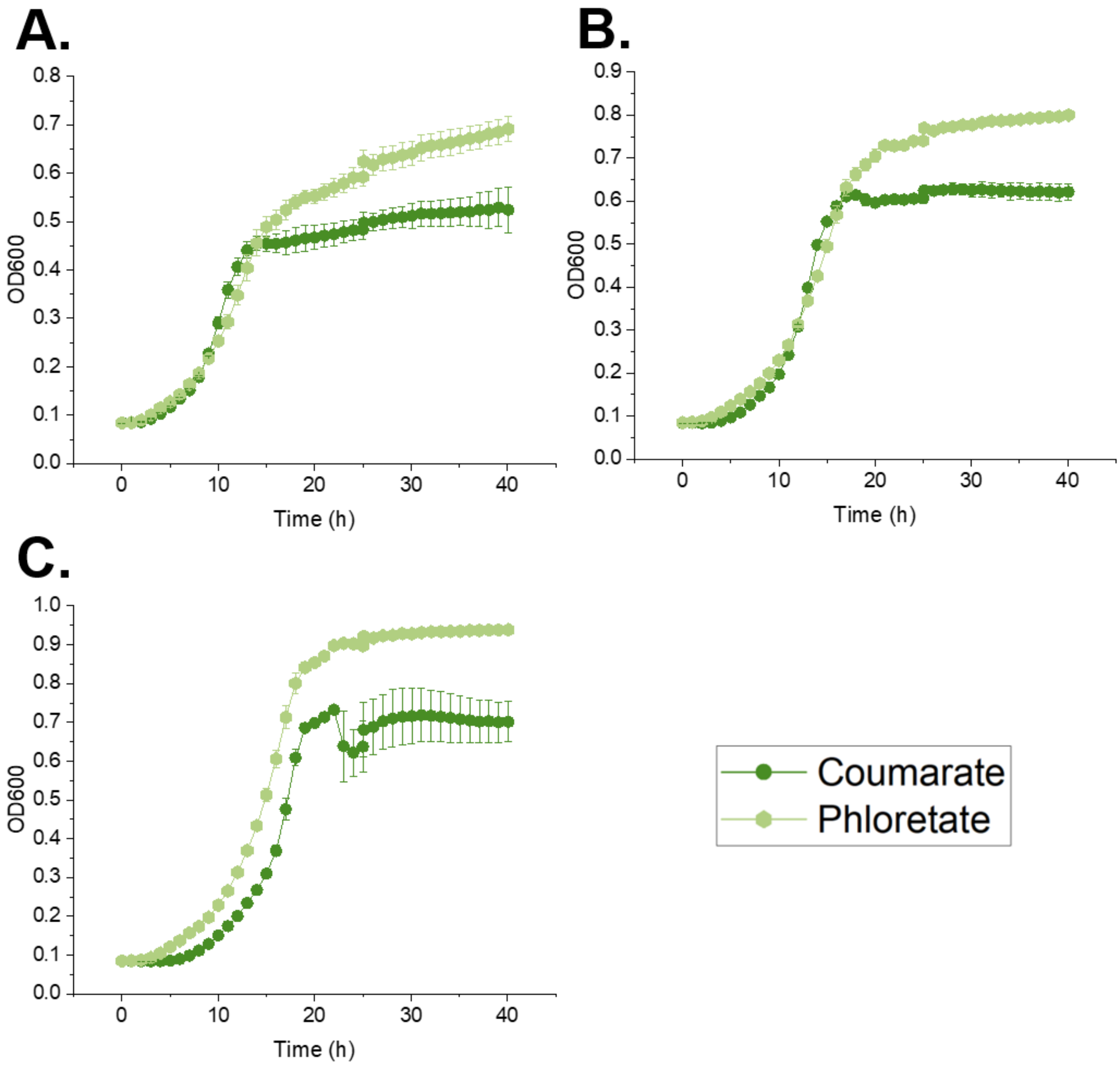


*Figure S3: The growth of ADP1 on coumarate or phloretate modelling the aromatic metabolites obtained when A. woodii is cultivated on coumarate. Used concentrations are A.) 5 mM, B.) 7.5 mM and C.) 10 mM. The error bars indicate the standard deviations from the average values of the biologically independent triplicates. In some cases, the error bars are smaller than the size of the marker.*

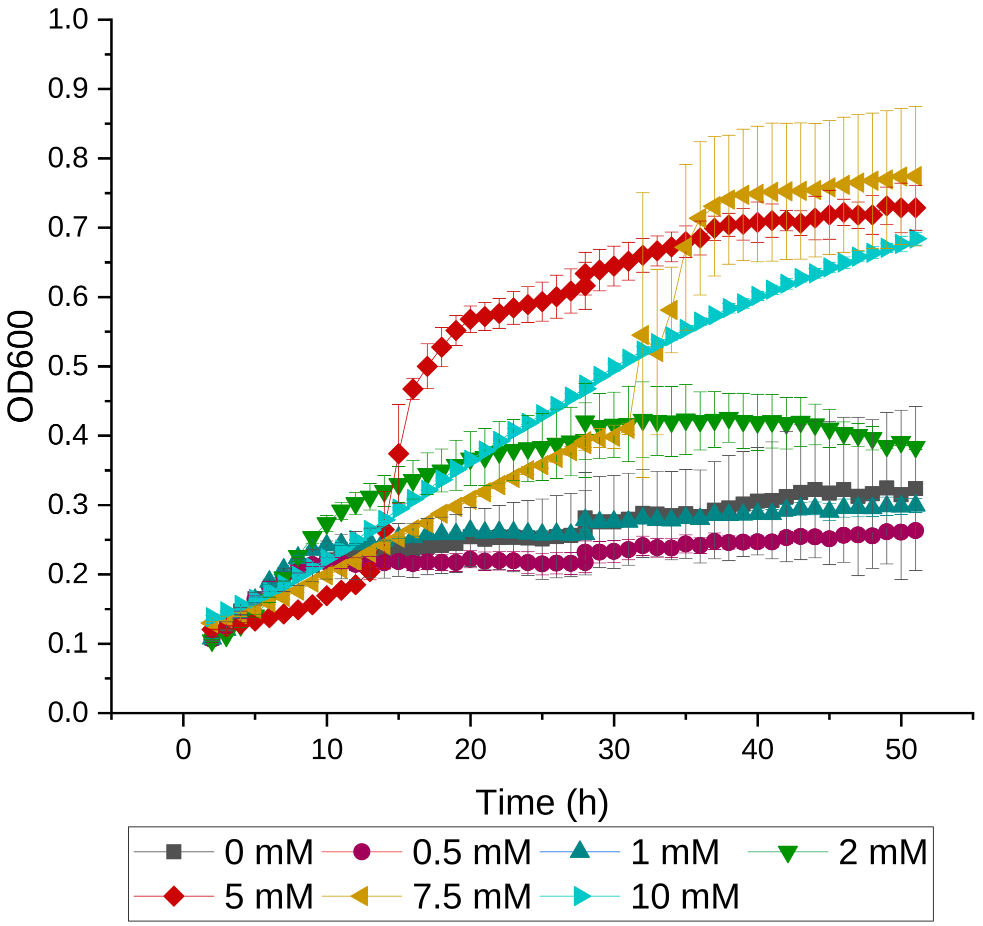


*Fig S4: The growth of ADP1 in MSM with catechol as a sole carbon source. The cultivation volume was 200 µl. The linear increase of OD in 10 mM cultivations was caused by the color change of catechol in the medium and therefore it is not representing the accumulation of biomass. The error bars indicate the standard deviations from the average values of the biologically independent triplicates. In some cases, the error bars are smaller than the size of the marker*

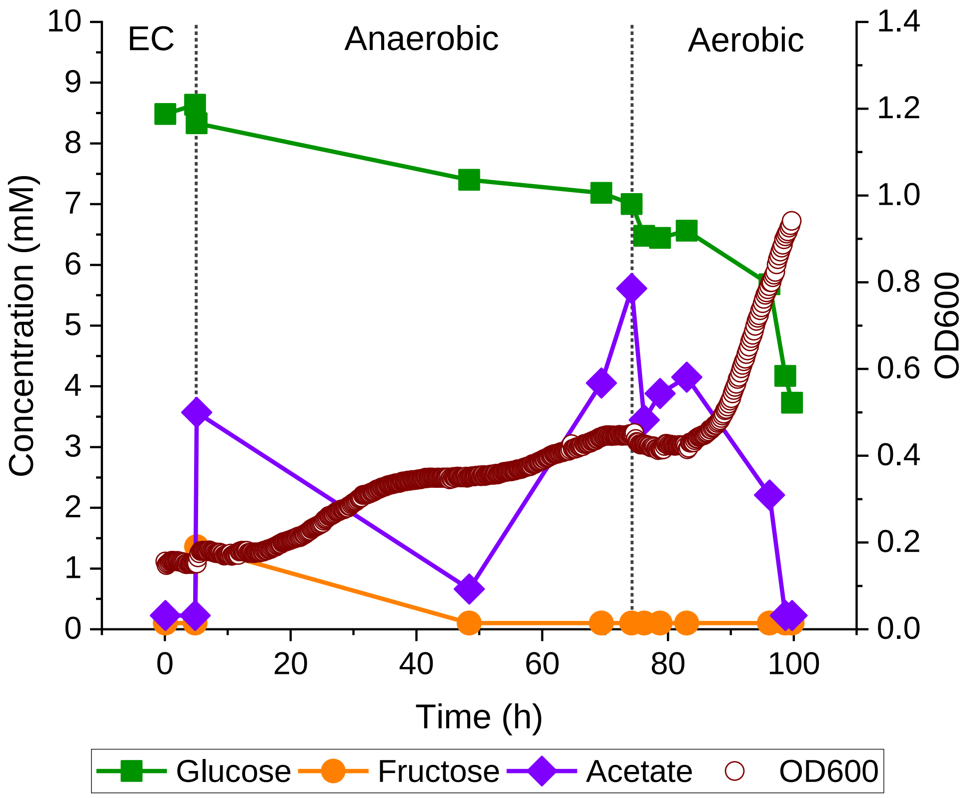


*Figure S5: Experimental data of the three-phase one-pot coculture of A. woodii and ADP1ΔcatBC. In this graph, only the utilization and production of sugars and organic acids are presented. Glucose is supplemented in the medium at the beginning of the cultivation. Fructose and acetate peaks at the first dashed lined are caused by the inoculation of A. woodii. Please notice that compared to Fig. 5, the presented acetate concentration here describes the overall concentration of acetate, not only the acetate produced by A. woodii.*
